# Inter-subunit coupling enables fast CO_2_-fixation by reductive carboxylases

**DOI:** 10.1101/607101

**Authors:** Hasan DeMirci, Yash Rao, Gabriele M. Stoffel, Bastian Vögeli, Kristina Schell, Aharon Gomez, Alexander Batyuk, Cornelius Gati, Raymond G. Sierra, Mark S. Hunter, E. Han Dao, Halil I. Ciftci, Brandon Hayes, Fredric Poitevin, Po-Nan Li, Manat Kaur, Kensuke Tono, David Adrian Saez, Samuel Deutsch, Yasuo Yoshikuni, Helmut Grubmüller, Tobias J. Erb, Esteban Vöhringer-Martinez, Soichi Wakatsuki

## Abstract

Enoyl-CoA carboxylases/reductases (ECRs) belong to the most efficient CO_2_-fixing enzymes described to date. However, the molecular mechanisms underlying ECR’s extraordinary catalytic activity on the level of the protein assembly remain elusive. Here we used a combination of ambient temperature X-ray Free Electron Laser (XFEL) and cryogenic synchrotron experiments to study the structural organization of the ECR from *Kitasatospora setae. K. setae* ECR is a homo-tetramer that differentiates into a dimer of dimers of open- and closed-form subunits in the catalytically active state. Using molecular dynamics simulations and structure-based mutagenesis, we show that catalysis is synchronized in *K. setae* ECR across the pair of two dimers. This conformational coupling of catalytic domains is conferred by individual amino acids to achieve high CO_2_-fixation rates. Our results provide unprecedented insights into the dynamic organization and synchronized inter- and intra-subunit communications of this remarkably efficient CO_2_-fixing enzyme during catalysis.

**Significance Statement:** Fixation of CO_2_ offers real potential for reaching negative CO_2_ emissions in bioenergy, and bioproduct utilization. The capture and conversion of atmospheric CO_2_ remains a challenging task. Existing biological systems can be exploited and optimized for this use. Bacterial enoyl-CoA carboxylases/reductases (ECRs) encompass the fastest CO_2_-fixing enzymes found in nature to date. However, the mechanisms underlying ECR’s extraordinary catalytic activity remain elusive. Our structural, computational, and biochemical results elucidate the dynamic structural organization of the ECR complex and describe how coupled motions of catalytic domains in the ECR tetramer drive carboxylation. This mechanistic understanding is critical for engineering highly efficient CO_2_-fixing biocatalysts for bioenergy and bioproduct applications.

---

The capture and conversion of atmospheric CO_2_ remains a challenging task for chemistry, resulting in an ever-increasing interest to understand and exploit CO_2_ fixation mechanisms offered by biology^1^. The family of enoyl-CoA carboxylases/reductases (ECRs) encompasses the most efficient CO_2_-fixing enzymes found in nature to date. In respect to catalytic rate, ECR family members from primary metabolism outcompete RubisCO, the central enzyme in photosynthesis, by more than one order of magnitude, which makes them exquisite models to study the molecular base of efficient CO_2_ catalysis ^2,3^.

ECRs catalyze the reduction of various α,β-unsaturated enoyl-CoAs using the reduced form of the cofactor nicotinamide adenine dinucleotide phosphate (NADPH). Hydride transfer from NADPH to the substrate generates a reactive enolate species, which acts as a nucleophile that attacks a bound CO_2_ molecule (**Figure 1a**)^2,3, 4,5^. However, the structural details of the carboxylation reaction have remained elusive, in part due to the lack of high-resolution structures of ECRs containing catalytic intermediates and carboxylated products.

**Figure 1:**
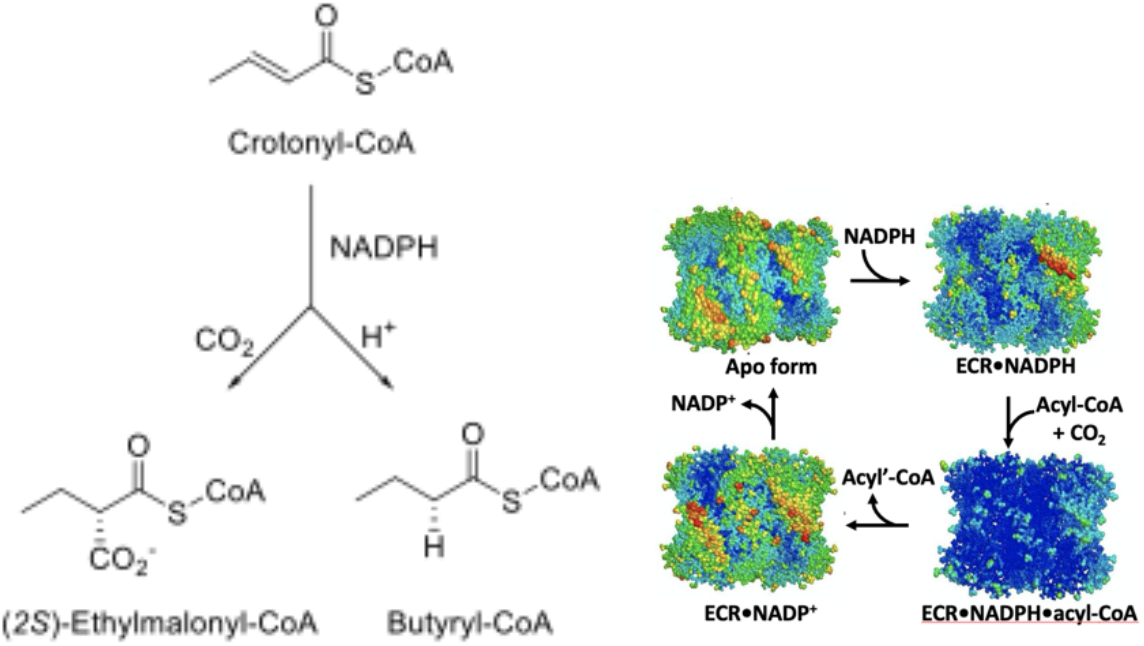
Reaction scheme and structural organization of the K. setae ECR complex. **a**. Carboxylation reaction scheme of ECR. **b**. Anisotropic B-factors of the tetramer of the different ECR complexes solved in this study are shown color coded according to the B factors (blue low and red high values).

Until recently, five structures of ECRs have been reported. These ECRs all have different substrate specificities, ranging from short- (PDB: 3HZZ, 3KRT) to long-chain (4A0S^6^) and aromatic enoyl-CoA substrates (4Y0K^7^), and are from different biological backgrounds, including primary (i.e., central carbon) metabolism (PDB: 4GI2) and secondary metabolism (PDB: 4A0S, 4Y0K). However, most of them were co-crystalized with NADPH or NADP^+^ only and do not contain CO_2_, enoyl-CoA substrates, or acyl-CoA products.

In a recent study, we reported the ternary structure of ECR (EC 1.3.1.85) from *K. setae* in complex with ethylmalonyl-CoA and NADPH (6OWE) and identified four active site residues critical to binding and accommodation of CO_2_ at the monomer level^8^. Yet, the molecular mechanism of the extraordinary rate enhancement of CO_2_ fixation by ECR tetramers remained enigmatic. This is even more notable, as ECR complexes from primary metabolism, such as *K. setae* ECR, show on average almost 30-fold higher CO_2_ fixation rates compared to those ECR complexes from secondary metabolism, despite very high structural similarities. Overall, this work indicated that ECRs from primary metabolism must use specific mechanisms that allow them to enhance CO_2_ fixation dramatically compared to their structural closely related homologs from secondary metabolism.

Here, we aimed at providing a mechanistic understanding of the fast carboxylation reaction of *K. setae* ECR on the level of the overall oligomeric complex. We first determined four high-resolution *K. setae* ECR structures in different conformational states: the apo form and three holo forms in a binary complex with the reduced cofactor NADPH, in a ternary complex with NADPH and butyryl-CoA, and a binary complex with the oxidized cofactor NADP^+^ (**Figure 1b**).

Our high-resolution structures show that the tetrameric complex assumes a dimer-of-dimers geometry (“a pair of dimers”). In this complex, the central oligomerization domains of *K. setae* ECR remain essentially unchanged, while the peripheral catalytic domains move significantly to provide two sets of active site conformations – open- and closed-form – upon binding of the NADPH cofactors and the presence of substrates. This coordinated motion is enabled by a tight coupling of catalytic domains across the pairs of dimers, which we could trace down to the role of individual amino acids far away from the active site. Using computer simulations and kinetic experiments, we provide compelling evidence that synchronization across the pair of dimers through “swing-and-twist-motions” is crucial for *K. setae* ECR to achieve high catalytic rates. We further demonstrate that subunit communication within a dimer is essential to synchronize open- and closed states. Altogether, our results unveil a dynamic inter-subunit synchronization of the *K. setae* ECR complex that is essential for the functional organization of one of nature’s most efficient CO_2_-fixing enzymes during catalysis.

## RESULTS

### Apo ECR is a symmetric homotetramer, readily accessible for cofactor and substrate binding

We first determined the apo form of the ECR crystal structure from *K. setae* at 1.8 Å resolution using synchrotron X-ray crystallography at cryogenic temperature (**SI, Table S1**). The asymmetric unit contains one homotetramer composed of four subunits arranged in a dimer of dimers geometry (“pair of dimers”) similar to those of the previously reported binary and ternary ECR complexes, crotonyl-CoA carboxylase/reductase AntE in complex with NADP^+^, (PDB: 4Y0K) and 2-Octenoyl-CoA carboxylase/reductase CinF in complex with NADP+ and 2-Octenoyl-CoA, (PDB: 4A0S), respectively. Overall, the *K. setae* ECR tetramer shows a non-crystallographic, close to *D*_2_ (dihedral) symmetry with four conformationally identical subunits (**SI, Figures S1-S2**). The tetrameric oligomer state of the ECR apo enzyme is further supported by size-exclusion chromatography which showed that the 51.2 kDa protein eluted as a single peak at 205 kDa (**SI, Figure S3**).

Each ECR subunit consists of two domains – a larger catalytic domain formed by residues 1-212 and 364-445, and a smaller oligomerization domain formed by residues 212 to 363 (**SI, Figure S4A**). The oligomerization domain comprises a Rossmann fold^9^ with repeating αβ-motifs that form a 6-stranded β-sheet (β12 to β17). The 6-stranded β-sheets of two neighboring subunits are combined into one 12-stranded β-sheet, forming the core of one dimer, subunits A/C or B/D. Two of these Rossmann fold domains form the tetrameric complex’s core (**SI, Figure S4B**). The catalytic domains of *K. setae* ECR are located at the periphery of the tetrameric complex. The active site cavities in the apo form are open and accessible for both the cofactors and substrates.

### Cofactor binding breaks tetramer symmetry and induces dimer-of-dimers formation

We first analyzed how cofactor binding affects the enzyme by determining the *K. setae* ECR-NADPH binary complex’s crystal structure at 2.4 Å resolution using serial femtosecond X-ray crystallography (SFX) at ambient temperature (**Figure 2, Table S1**)^10–13^. In all four subunits, NADPH binds with its adenine moiety in the oligomerization domain and spans the catalytic domain, where its nicotinamide moiety is located (**SI, Figure S5**).

**Figure 2:**
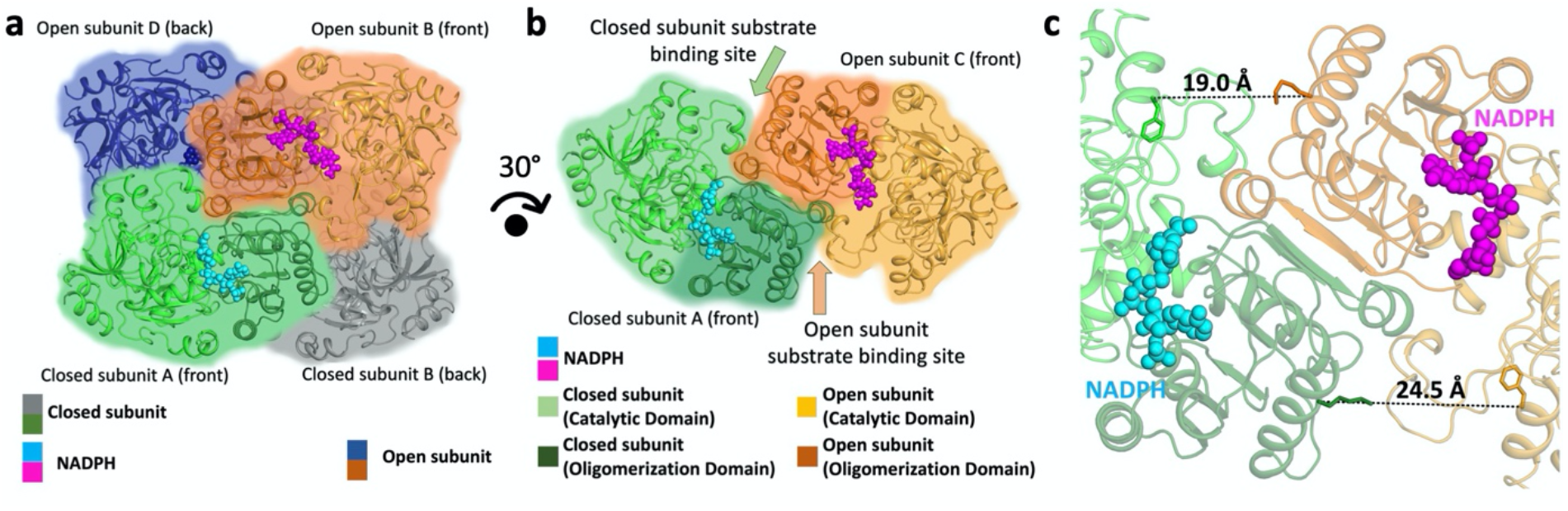
Binding of NADPH results in global and local conformational changes in K. setae ECR. **a**. NADPH bound tetramer complex that is organized as dimer of dimers, a pair of closed (green) and open (orange) subunits and another pair containing closed (gray) and open (blue) subunits. **b**. The foreground dimer with open- (orange) and closed-form (green) subunits rotated by 30 degrees from the view in Figure 2a. Each subunit is composed of a catalytic and an oligomerization domain. **c**. Comparison of the putative substrate binding sites between the open and closed-form subunits.

Notably, NADPH binding breaks the dihedral *D*_2_ symmetry observed in the apo-form tetramer, while symmetry about the y-axis is retained, resulting in a non-crystallographic, almost cyclic *C*_2_ symmetry (**SI, Figure S1**). In the NADPH•ECR binary complex, each of the two dimers, A/C and B/D, loses its 2-fold symmetry, giving rise to closed (A and B) and open (C and D) subunits (**Figure 2a-b and Figure S1**). In the A and B subunits, the cofactor binding pocket is compressed inwards, which seals the NADPH cofactor within the catalytic domain, resulting in a “closed-form” state (**Figure 2b**). On the other hand, the C and D subunits show open cofactor-binding pockets, referred hereafter as an “open-form” state. The substrate-binding pocket in the open-form C or D subunit is more than 5 Å wider than the closed A or B subunit (**Figure 2c**). Taken together, these structures indicated that during catalysis, the enzyme differentiates into a dimer of dimers (A/C and B/D dimers, respectively), with one open (A or B) and one closed subunit (C or D) per dimer.

### Cofactor-substrate complex indicates half-site reactivity within each dimer

We next investigated the effects of substrate binding to the *K. setae* ECR tetramer complex and determined its ternary complex structure with the substrate analog butyryl-CoA and NADPH at 1.7 Å resolution (**Figure 3**). The structure is overall very similar to the conformation of the ECR•NADPH binary complex with the non-crystallographic, pseudo *C*_2_ cyclic symmetry (**Figure 3a** and **Figure S1**). It comprises two sets of open- and closed-form subunits that overlay very well with the ones of the ECR•NADPH binary complex (**SI, Figures S1-S2**). The NADPH cofactor appears bound to all active sites; however, only the closed-form subunits A and B also contain electron density for the complete butyryl-CoA thioester, while the open-form subunits (C and D) show only the adenine group electron density of the butyryl-CoA thioester (**Figure 3a-b**). The presence of the intact butyryl-CoA thioester in the active site of the closed-form subunits strongly suggests that the A and B conformation represent the Michaelis complex in which substrate and cofactor are positioned for catalysis. In contrast, the open-form subunits (C and D) represent catalytically incompetent ternary complexes.

**Figure 3:**
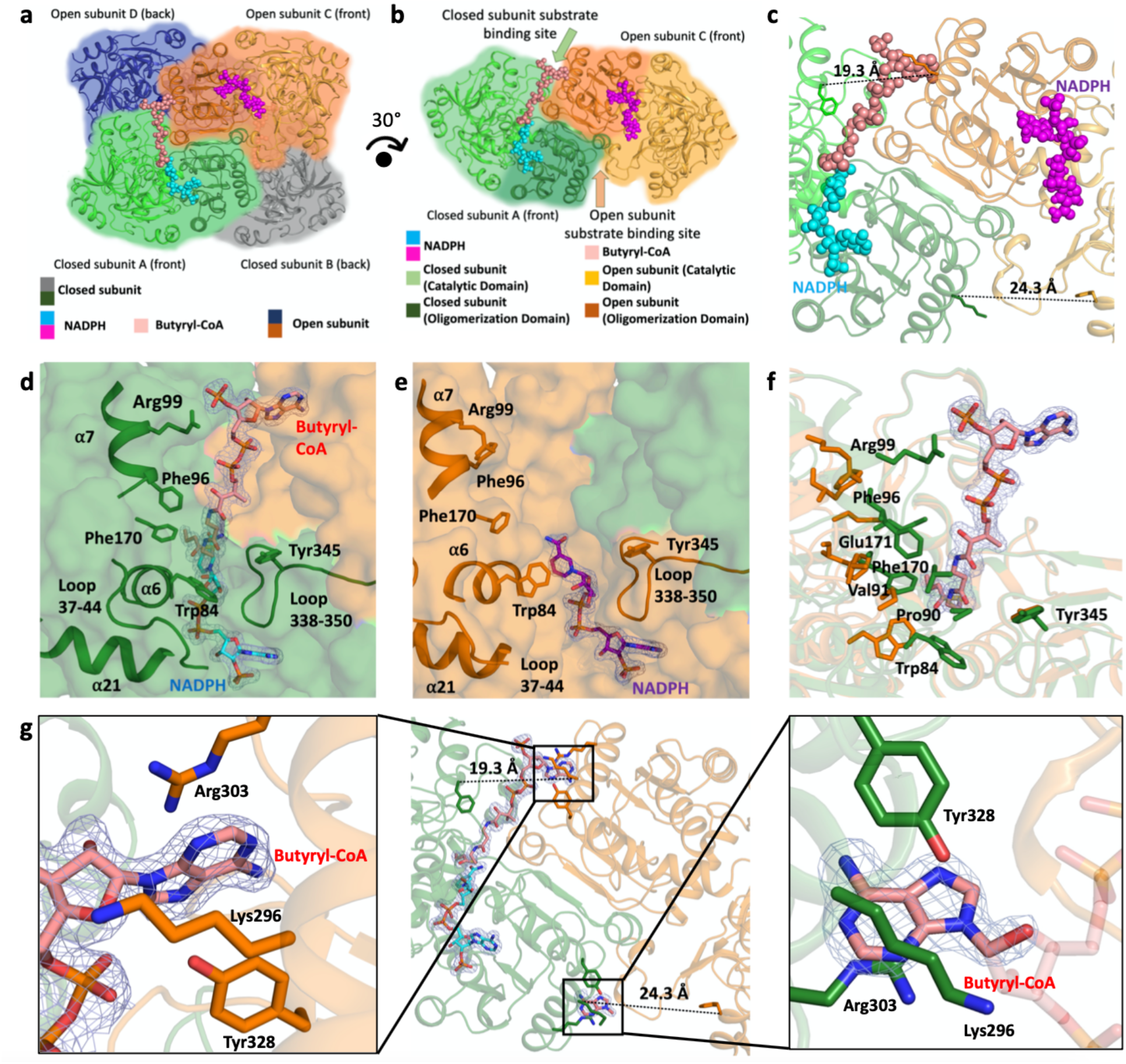
Structure of the ternary ECR complex. **a**. ECR tetramer in complex with NADPH and butyryl-CoA organized as dimer of dimers, the foreground dimer with one closed subunit (green) with NADPH and butyryl-CoA and open (orange) subunit containing NADPH, and another pair in the background with one closed (gray) and open (blue) subunits. **b.** The foreground dimer with closed (green) and open (orange) subunits, rotated by 30 degrees from the view in Figure 3a. Butyryl-CoA and NADPH atoms are represented as spheres. **c**. Comparison of the product binding site between the open and closed-form subunits. **d**. Cartoon and stick representation of the closed-form subunit active site. In panels **d** to **g**, simple 2F_o_-F_c_ density contoured at 1.5 σ level is shown for butyryl-CoA, or portion thereof, and NADPH within 3 Å from the molecules. **e**. Cartoon and stick representation of the open-form subunit active site. **f**. Superposition of the open-form subunit onto the closed-form subunit with stick representation of the residues surrounding butyryl-CoA. **g**. Butyryl-CoA binding in both open and closed-form subunits with electron density of the bound butyryl-CoA and NADPH at the active site of the closed subunit (green) and the adenine ring of butyryl-CoA at the active site of the open subunit (only the adenine ring electron density is visible). Left inset: the adenine binding pocket of the open-form subunit stabilizing the adenine ring of butyryl-CoA that stretches into the adjacent closed-form subunit. Right inset: the adenine binding pocket of the closed-form subunit holding the adenine ring of butyryl-CoA. Note that only the adenine ring of butyryl-CoA is visible while the rest of the molecule is disordered, hence indicated in transparent stick model. In both cases, three residues of the adjacent subunits, Lys296, Arg303, and Tyr328 together hold the adenine ring.

Binding of cofactor and substrates in the closed-form ECR subunit is achieved by loops 37-44, 88-94, 338-350, and helices 6, 7, and 21 of the catalytic domains, which creates multiple interactions of the protein with the NADPH and butyryl-CoA (**Figure 3d-f and Figure S11**). Notably, the CoA-thioester extends from the closed-form active site out to the oligomerization domain of the neighboring open-form subunit in the same dimer pair (from A to C in A/C dimer, or from B to D in B/D), where Arg352 and Tyr353 from the open-form subunit (C or D) interact with the phosphate backbone of the CoA. The adenosine tail of the CoA extending from the closed-form subunit interacts with three residues Tyr328, Lys296, and Arg303 that form an adenine binding pocket at the neighboring open-form subunit (**Figure 3g**). When we inspected CoA-thioester binding in the open-form subunits, we also observed the electron density in the adenine binding pocket of the neighboring closed form subunits (A or B), strongly indicating that the CoA-thioester was bound (**Figure 3g right inset**). However, the electron density beyond the adenine ring becomes invisible, suggesting that the part of the CoA molecule reaching into the open-form subunit’s active site remains flexible and disordered, which is corroborated by the higher B-factors of the catalytic domain (**Figure 1B bottom right**).

To evaluate the substrate’s flexibility in the open- and closed-form subunits, quantum mechanical/molecular mechanics (QM/MM) simulations on a dimer of subunits A and C were performed. These simulations showed that the substrate in the closed subunit had significantly lower temperature factors (B-factors) than the one in the open subunit (**SI, Figure S6-7, movies S1a-e, S2a-e**). In the open subunit, the acyl moiety shows a high degree of flexibility in the active site in agreement with the higher B factors observed in crystal structures.

Together, crystallographic structures and QM/MM simulations indicate that upon substrate binding the enzyme complex splits into two catalytically competent (A or B) and incompetent subunits (C or D) per dimer, suggesting that the enzyme tetramer operates with half-site reactivity, in which catalytically competent and incompetent active sites alternate during catalysis^15–17^.

### Swing and twist motion of peripheral catalytic domains are coupled within and across pair of dimers to promote catalysis

To study the switching between the open- and closed-forms of the ternary complex, we used computer simulations. Starting from the ternary crystal structure with NADPH and butyryl-CoA, we modeled the natural substrate crotonyl-CoA to carry out all-atom molecular dynamics simulations of the catalytic active enzyme (see SI Methods). In our simulations, the closed subunit remained closed and maintained crotonyl-CoA in the active site when the dynamics of the substrate were restrained to maintain specific protein-substrate/cofactor interactions. However, in simulations without substrate restraints, the acyl moiety of the substrate left the active site resulting in a conformational change of the closed subunit to the open form.

To study this conformational change in detail, we eliminated the crotonyl-CoA substrate from the ternary complex creating an ECR•NADPH binary complex with two closed and two open subunits. In all ten 100ns simulations of the tetramers, the closed subunits underwent a fast conformational change during the first tens of nanoseconds (the same was observed when we started from the X-ray binary ECR•NADPH structure). To further describe this conformational change, we used Principal Component Analysis (PCA) of the backbone Ca-atom’s position of ten combined 100 ns trajectories of both dimers. The first principal component (PC1) represents transition of the closed to the open form, and the second principal component (PC2) describes a twist motion of one subunit to the other subunit in each dimer. In both cases the catalytic domains move relatively to the oligomerization domains (**Figure 4a-c and SI Movies S3-S4**). The rotation axes of PC1 and PC2 cross with each other with close to 90 degrees inside the catalytic domain in each subunit. The two PC1 axes of subunits A and C are nearly parallel, corresponding to the closed-open motions, while the PC2 axes show the twisting motions of the catalytic domains (**Figure 4b-c**).

**Figure 4:**
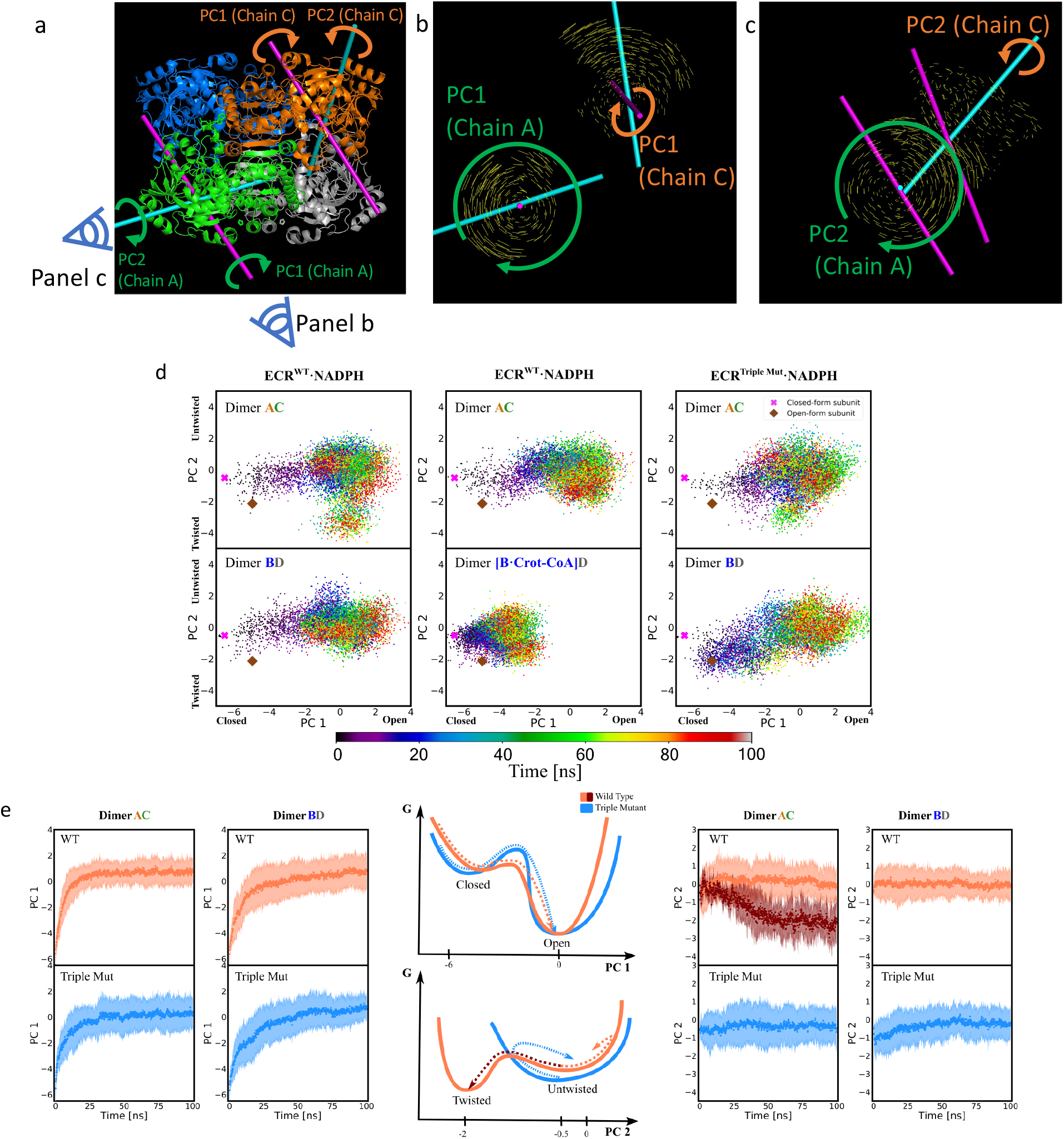
Molecular dynamics simulations and Principal Component Analysis reveal that absence of substrate induces transition between open and closed forms of the catalytic domain in the ECR tetramer and a twist in the subunits, which would move a bound substrate towards the NADPH cofactor. **a**. Representation of the swing motion of the catalytic domain in dimer AC described by the first principal component (PC1) and the twist motion of one subunit to the other by PC2. This twist motion would bring a bound substrate closer to the NADPH cofactor. PC1 and PC2 axes of A (closed) and C (open) subunits are superimposed on the ternary tetramer structure in the same orientation as in Fig. 3A, showing the functional relevance of PC1 as opening-closing and PC2 as twisting of the catalytic domains. **b** and **c**. Sets of vectors between initial and final Ca positions of the catalytic domains are shown viewing into the PC1 axis (panel **b**) and the PC2 (panel **c**) axes of subunit A as shown on the bottom left of panel **a**. **d**. Projection of ten 100ns trajectories on the first two principal components of the wild type binary system with NADPH cofactor ECR^wt^·NADPH (cross and diamond represent the open- and closed form subunit from the X-Ray structure). The middle panels show the projection on the same PCs for the ECR^wt^ ternary complex bearing only one substrate in the B subunit of the BD dimer ([B·Crot-CoA]D). The right panels represent the dynamics of the E151D/N157E/N218E triple variant with the NADPH cofactor (ECR^Triple.Mut^·NADPH) on the same eigenvectors. Colors of each points represent time frame according to scale bar at the bottom. **e**. Mean values and 1.5 times the standard deviation of the principal components of ten trajectories as function of simulation time for each dimer in the WT (orange) and E151D/N157E/N218E triple variant (blue) ECR·NADPH complex. Time traces of PC1 are shown on the left and of PC2 on the right. In the middle, free energy profiles as function of PC1 and PC2 were derived for the wild type (orange) and variant (blue) based on the observed kinetics. The triple variant with slower opening kinetics (PC1) is consistent with a barrier in the transition to the open form and its larger standard deviation suggest a shallower minimum in the closed form. The free energy profile of PC2 corresponds to a broad minimum for the triple variant because of the larger standard deviation observed in the kinetics and a large barrier to reach the twisted conformation observed in the WT dynamics (brown trajectory shown in dimer AC for PC2 on the right).

To characterize the dynamics, we compared the projections of all ten dimer trajectories on PC1 and PC2. Dimer A/C opens its closed subunit first, associated with PC1, followed by a twist motion characterized by PC2 in some of the simulations (**Figure 4d and SI Figure S9**). This twist motion rotates the catalytic domain of the empty, now open form active site in subunit A, which would push a bound crotonyl-CoA substrate towards the NADPH cofactor in the C subunit. The B/D dimer’s dynamics on the other hand were mainly governed by the closed subunit opening represented by PC1, likely because of initial structural differences of the two dimers.

After characterizing the conformational transitions associated with the presence or absence of CoA substrate, we asked whether the dynamics of one dimer in the tetramer complex is coupled to the other dimer. Computer simulations allowed us to address this question by studying the dynamics of a hypothetical system that lacks the substrate in only one active site. Removing crotonyl-CoA from subunit A, we expected the – now empty – “closed” active site subunit to transition to the open form inducing the twist motion in the A/C dimer observed before, and the other closed subunit B still bearing the substrate to remain closed, if the two dynamics of the pair of dimers are uncoupled. The projection of ten 100 ns trajectories on PC1 and PC2 confirmed that the empty active site (A in the A/C dimer) undergoes the conformational change to the open form (**Figure 4d**), whereas the dimer containing the substrate (B in the B/D dimer) remained in a closed conformation (see **SI Figure S8** for comparison to the simulation with two substrates maintaining the closed state). Importantly, however, none of the A/C dimer trajectories showed a significant change in PC2 associated with the twist motion observed in the binary ECR•NADPH complex, indicating that the twist motion is only initiated once the other dimer B/D adopts the open conformation.

In summary, our molecular dynamics simulations indicate that absence (or release) of the substrate triggers conformational transition from the closed to the open conformation and that coupled dynamics of the two dimer pairs induce an (additional) twist motion after opening the closed subunit of the other dimer pair.

### Three remote amino acids couple catalysis between pairs of dimers

Given the coordinated motions of the catalytic domains during catalysis, how is catalysis synchronized across the enzyme complex (i.e., across dimers A/C and B/D)? One intriguing aspect of the ECR tetramer structures is that the catalytic domains share a common interface of 1636 Å^2^ between the pairs of dimers (**Figure 5a-b**), suggesting that they move together as rigid bodies.

**Figure 5.**
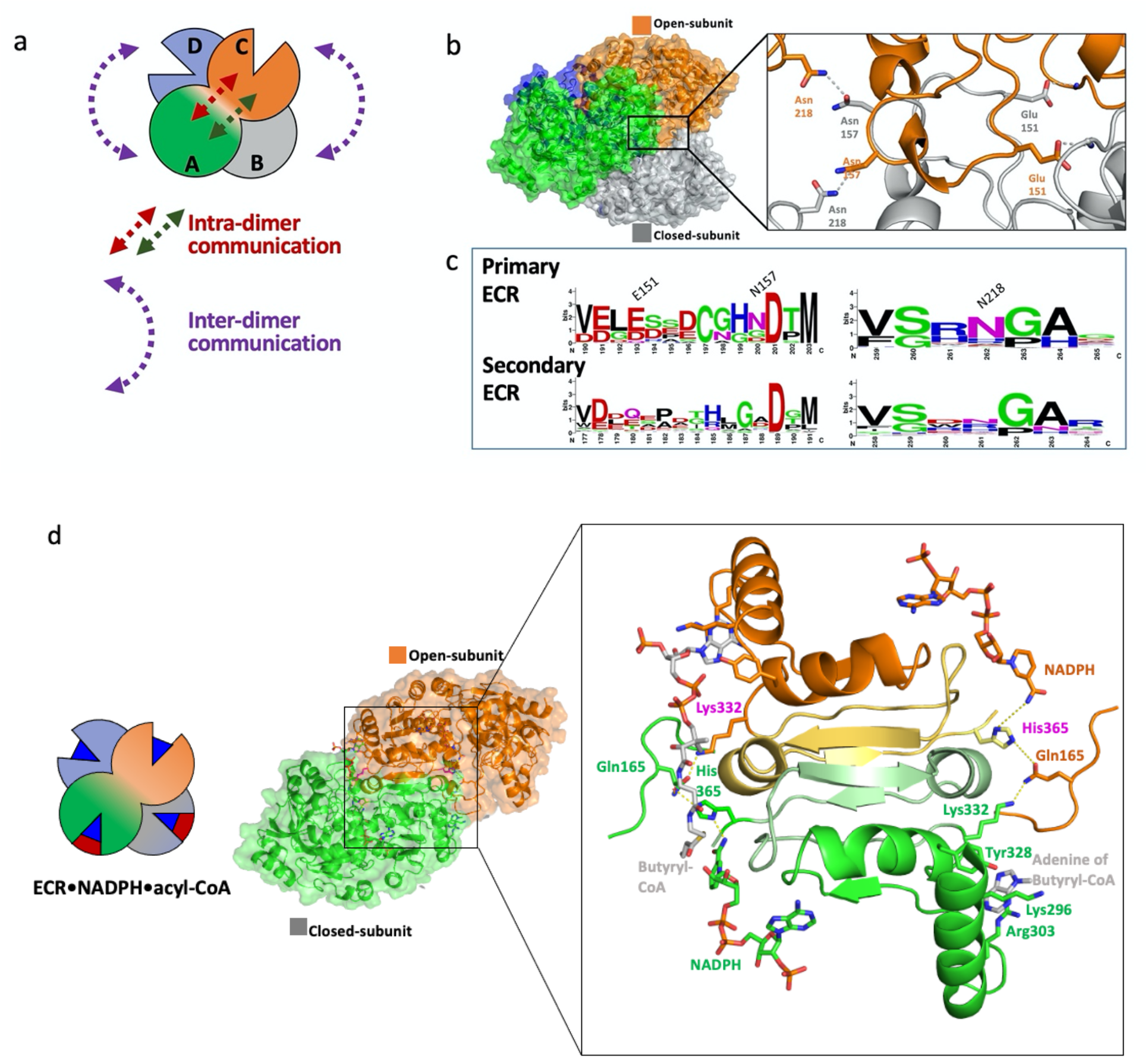
Inter- and intra-dimer communications drive fast CO_2_ fixation by K. setae ECR. **a.** Two distinct sets of communications: inter-dimer interactions between the catalytic domains from two dimers (purple arrows) and intradimer communication between the open and closed subunits within each dimer (brown arrows). **b**. Inter-dimer catalytic domain interface and positions of selected amino acids that were mutated in this study to affect the interface between the two catalytic domains (open-form subunit in orange and closed form subunit in gray). The right panel shows the mutual H-bonding interaction between Asn218 and Asn157 from open and closed form subunits and H-bonding between Glu151 and N-atom from protein backbone. **c**. Alignment of ECR protein sequences from the primary (upper row) and the secondary (lower row) metabolism represented as sequence logos. Numbering of residues, above first row, is according to their position in K. setae ECR. **d**. Communication between the closed (green) and open (orange) subunits across the two dimers of K. setae ECR. In the closed conformation the contacts between NADPH-His365-Glu165 and Lys332 of the adjacent open subunit allow for the correct intra-dimer communication. In the open conformation the communication network is compromised as indicated by the increased distances between the amino acid sidechains that cause the incorrect positioning of the nicotinamide ring of NADPH.

What are the molecular determinants that synchronize catalysis across the pair of open/closed form dimers? The inter-catalytic domain interface is predominantly hydrophobic with some electrostatic interactions (**Figure 5b**). Most notable are Asn218, which forms a hydrogen bond to Asn157 of the adjacent subunit of the other dimer, and Glu151 interacting with the backbone nitrogen atom of Asn133 (and/or Ala134) of the neighboring subunit of the other dimer (**Figure 5b**). Multiple sequence alignment showed that Glu151, Asn218, and Asn157 are highly conserved in ECRs from primary (i.e., central carbon) metabolism with faster CO_2_-fixation kinetics (average *k_cat_* 28 s^-1^), but not in ECRs from secondary metabolism with average *k_cat_*=1.2 s^-1^ (**Figure 5c**), which raised the question about their potential role in promoting catalysis in primary metabolic ECRs.

Notably, mutation of these residues that are more than 20 Å away from the active site dramatically affected the kinetic parameters of *K. setae* ECR (**Table 1**). In the E151D variant, the *k_cat_* value decreased by a factor of five, demonstrating that weakening the interaction between catalytic domains has profound effects on the enzyme’s catalytic rate. Mutations that targeted the asparagine interaction network also showed substantial effects on the catalytic rate and affected *K_M_* of substrate binding. Most notable were variants N218E single and E151D/N157E/N218E triple variants that decreased the *k_cat_* by more than 25- and 100-fold, respectively, highlighting that communication through the interface of the catalytic domains of the pair of dimers is an essential determinant of the catalytic rate in *K. setae* ECR.

**Table 1.** Steady state analysis of *K. setae* ECR and variants targeting the catalytic domain interface between the pair of dimers. (Michaelis-Menten curves of *K. setae* ECR and its variants are provided in **SI, Figure S12**)

|  | Crotonyl-CoA |  |  | NADPH |  | CO<sub>2</sub> |  |
| --- | --- | --- | --- | --- | --- | --- | --- |
| Enzyme | K<sub>M</sub> (μM) | K<sub>i</sub> (μM) | k<sub>cat</sub> (s<sup>-1</sup>) | K<sub>M</sub> (μM) | k<sub>cat</sub> (s<sup>-1</sup>) | K<sub>M</sub> (μM) | k<sub>cat</sub> (s<sup>-1</sup>) |
| Wild type | 21 ± 2 | 3650 ± 810 | 103 ± 3 | 37 ± 4 | 86 ± 2 | 90 ± 10 | 78 ± 2 |
| E151D | 28 ± 2 | 1960 ± 250 | 20 ± 1 | 72 ± 11 | 17 ± 1 | 80 ± 10 | 21 ± 1 |
| N157E | 515 ± 75 | - | 22 ± 1 | 105 ± 27 | 17.1 ± 0.4 | 40 ± 6 | 14 ± 0 |
| N218E | 272 ± 37 | - | 3.7 ± 0.2 | 66 ± 9 | 3.3 ± 0.0.2 | 840 ± 150 | 3.3 ± 0.2 |
| E151DN157EN218E | 245 ± 25 | - | 1.1 ± 0.0 | 26 ± 3 | 0.7 ± 0.0 | 440 ± 40 | 0.9 ± 0.0 |

To exclude that the oligomerization state of the different complexes was not altered through these mutations, we used gel filtration and native gel analysis (**SI Figure S3**). Gel filtration under the same conditions as our kinetic measurements showed that all enzyme variants remained tetramers, while native gel analysis conducted under more disruptive conditions showed slight increases in the dimer and monomer fractions. Overall, our mutational and kinetic data supported the hypothesis that synchronization of catalytic domains contributes to the catalytic rate and is conferred through the hydrogen-bond network at the pair of dimers’ interfaces.

To study the effect of the E151D/N157E/N218E mutations on the synchronization of catalytic domains and the conformational transitions of the binary NADPH•ECR complex we used computer simulations as discussed above. Projection of ten 100ns trajectories on the previously defined eigenvectors displayed the same opening transition of both dimers in the initially closed binary NADPH•ECR triple variant complex (Figure **4d**). We analyzed the time traces of their first two principal components (mean ± 1 std) of both dimers in the tetramer to spot the differences between the WT enzyme and the triple variant (see Figure **4e**). Notably, the triple variant opens the closed form slower than the WT enzyme and presents a broader distribution of values in both dimers during transition. The observed kinetics are consistent with a free energy profile along PC1 according to which the variant must cross a higher free energy barrier to reach the open form (scheme in **Figure 4e**). The broader distribution during the transition is caused by a shallower energy minimum of the closed form of the triple variant. For PC2, the dynamics for the WT and the triple variant also differ. The triple variant presents a broader distribution associated with a shallow free energy minimum and shifted minima causing the two systems to evolve in opposing directions at the beginning of the simulation. Interestingly, only the WT can pass the free energy barrier to a twisted state in some of the trajectories (shown in brown in **Figure 4e**). Overall, these results confirm that the E151D/N157E/N218E mutation at the dimer interfaces changes the free energy landscape of conformational transitions considerably. The communication between the dimers is reduced, as the triple variant does not share the WT’s concerted dynamics that are characterized first by an opening motion and then a twist of one subunit to the other. This twist motion of the open subunit will push bound substrate to the NADPH cofactor and might be directly coupled to the first step of the ECR reaction, i.e., the hydride transfer.

In summary, computer simulations together with kinetic data of the triple variant indicate that coupling the dynamics of the two dimers is conferred by three conserved amino acids in ECRs from primary metabolism (Glu151, Asn157, Asn218) and essential to reach fast CO_2_ fixation kinetics compared to their counterparts from secondary metabolism that lack those residues.

### CoA binding synchronizes open and closed subunits within a dimer

Besides inter-dimer domain interaction, the two subunits in each dimer bind different parts of the substrate in the open- and closed-form through the adenine binding pocket and the active site (intra-dimer interaction, **Figure 5a**). We hypothesized that this shared substrate-binding between neighboring subunits could also contribute to the mechanism of fast, synchronized catalysis.

To understand the role of substrate adenine binding in catalysis, we characterized the kinetics of different single, double, and triple variants of the adenine binding site (**Figure 3g** and **Table 2**). Mutations in the adenine binding pocket, particularly of Arg303, strongly increased the apparent *K_M_* of the CoA-substrate as expected and decreased the enzyme’s apparent *k_cat_* by a factor of two to three. Notably, when we used crotonyl-panthetheine, a truncated substrate lacking the adenosine moiety, a comparable decrease in *k_cat_* was also observed for the WT enzyme, suggesting that substrate interactions at the adenine binding pocket indeed contribute to catalysis. However, when we compared the kinetics of the WT enzyme with truncated substrate to a triple variant K296A/R303A/Y328F, in which adenosine binding was completely disturbed, the two variants behaved almost identical, strongly supporting the argument that binding of the adenine residue of the CoA substrate in the WT is important for high catalytic activity.

**Table 2.** Apparent Michaelis-Menten parameters of *K. setae* ECR and variants targeting the adenine binding pocket as mean values ± standard error. (Michaelis-Menten curves of *K. setae* ECR and its variants are provided in **SI, Figure S12**)

|  | Crotonyl-CoA |  |  | Crotonyl-Pantetheine |  |
| --- | --- | --- | --- | --- | --- |
| Enzyme | K<sub>M</sub> (μM) | K<sub>i</sub> (μM) | k<sub>cat</sub> (s<sup>-1</sup>) | K<sub>M</sub> (μM) | k<sub>cat</sub> (s<sup>-1</sup>) |
| Wild type | 21 ± 2 | 3650 ± 810 | 103 ± 3 | 8660 ± 530 | 37 ± 1 |
| K296A | 107 ± 11 | - | 68 ± 2 | - | - |
| Y328F | 11 ± 2 | 4671 ± 1693 | 80 ± 3 | - | - |
| K296A/Y328F | 190 ± 30 | - | 39 ± 2 | - | - |
| K296A/R303A/Y328F | 2180 ± 280 | - | 29 ± 2 | 7770 ± 110 | 42 ± 3 |
| K296A/R303K/Y328F | 830 ± 140 | - | 53 ± 2 | - | - |
| Q165A | 27 ± 4 | - | 56 ± 3 | 5630 ± 520 | 8.6 ± 0.3 |
| K332A | 450 ± 130 | - | 38 ± 7 | 2980 ± 510 | 0.3 ± 0.0 |

We noticed that the substrate adenine binding pocket is directly followed by a loop, which carries a lysine residue (Lys332) that interacts with the neighboring subunit’s active site. Lys332 from the open-form subunit engages in a hydrogen-bonding network with the nicotinamide of the NADPH cofactor bound to the closed-form subunit through Gln165 and His365 of the neighboring subunit (**Figure 5d**). These interactions are not observed in the active site of the open-form subunit (**Figure 5d**), raising the question of whether the hydrogen bonding network connected to the adenine binding pocket might be necessary for catalysis.

In variants K332A and Q165A, *k_cat_* was decreased by two-to three-fold, and for K332A, an additional increase in the apparent *K_M_* for crotonyl-CoA was observed, indicating some effect of these residues on catalytic rate (as well as substrate binding in case of Lys332) (**Table 2**). When we tested these variants again with the truncated substrate analog crotonyl-pantetheine, the catalytic activity in the Q165A variant was reduced by four-fold, while that of the K332A variant was reduced by more than two orders of magnitude compared to the WT with truncated substrate. More notably, the *K_M_* had not changed much between the WT and the two different variants (where it has even slightly improved), strongly suggesting that Gln165 and Lys332 are not involved in substrate accommodation, but rather synchronization of catalysis between open and closed subunits within the dimer.

In conclusion, adenine binding and the loop carrying the residues Lys332 and Q165 are necessary to synchronize catalysis between the two subunits within the dimer to drive fast CO_2_-fixation in *K. setae* ECR, providing the molecular basis of how catalysis might be synchronized between the open- and closed-form subunits within one dimer.

## Discussion

Our structural studies of *K. setae* ECR revealed unprecedented details on the functional organization of one of nature’s most efficient and fastest CO_2_-fixing enzyme. During catalysis, the enzyme complex differentiates into distinct functional subunits. The binding of the NADPH cofactor and substrates forces the homo-tetrameric apo enzyme into a dimer of dimers. Each dimer is constituted of open- and closed-form subunits. In the closed-form subunits, the NADPH cofactor and CoA substrate are aligned with each other, suggesting that this is the catalytically competent state. The open-form subunits bind the cofactor and the substrates’ adenine rings, but the rest of the acyl-CoA substrate remains flexible and invisible in the active site. Thus, the open-subunit active sites represent a catalytically incompetent state that is preorganized for a subsequent round of catalysis. Overall, this structural reorganization of the ECR complex strongly supports the idea that the enzyme operates with “half-site reactivity”. Catalysis is synchronized across the enzyme tetramer and alternates between the open and closed-form subunits to increase the overall catalytic efficiency of the complex^15–18^.

It is worth noting that “half-site reactivity “does not assume (or require) half-site occupancy. Indeed, all four subunits of the *K. setae* ECR tetramer ternary complex contain both NADPH and substrates (products). However, in the open form subunits, only electron density for the adenine groups of the CoA thioester is observed (Figure 3g). The “half-site reactivity”is also consistent with a negative, rather than positive, cooperativity, as almost all our kinetics data indicate.

To understand if the conventional half-site reactivity theory can be applied to the ECR tetramer case, we created a model for the enzyme kinetics assuming positive or negative cooperativity between the pairs of dimers. Instead of treating the four subunits as independent entities, we hypothesized that the two pairs of ECR dimers would cycle through the CO_2_-fixing reactions in synchrony, i.e., when one dimer switches from the open-close to the close-open states, the other dimer would do the opposite, close-open to open-close states. Using this model, we simulated the case with just one dimer with two subunits working in negative or positive cooperativity using the method described by Hill and Levitski 1980 (ref 18), (**SI, Figure S10**). For a thorough investigation of how cooperativity, positive or negative, enhances the overall reaction rate of *K. setae* ECR, we incorporated all the parameters in Hill and Levitski’s model in our simulation. It turned out that only negative cooperativity could explain our experimental observations that ECR activity in the WT tetramer is increased by two orders of magnitude compared to the triple variant E151D/N157E/N218E, in which synchronization is decoupled **(Table 1)**. Our simulations yielded a 90% probability that both active sites are occupied with substrates/products and NADPH/NADP^+^ (p22), while only 10% are half-occupied (p12+p21) (**SI, Figure S10**). Almost none of the dimers are empty (p11), which agrees with the occupancy of our experimentally determined structures of the ternary complex.

To synchronize half-site reactivity, interaction of the catalytic domains across and within the pairs of dimers is crucial. When the latter is perturbed in *K. setae* ECR (e.g., through triple mutant E151D/N157E/N218E or by disturbing the adenosine binding pocket), the catalytic rate of the enzyme is severely diminished. This observation is consistent with other theoretical and experimental data on half-site reactivity. Synchronization of distant catalytic subunits has been calculated to enhance the catalytic rate of enzymes by up to 20-fold^19,20^. Mutation of a single amino acid that couples the two catalytic sites of heptose isomerase GmhA reduced the catalytic rate to 6% of wild-type activity^21^. Perturbing the interaction network in *Escherichia coli* thymidylate synthase, another example of an enzyme with half-site reactivity ^22^, caused a 400-fold decrease in *k_cat_*^23,24^, demonstrating that domain interactions are essential in promoting enzyme catalysis^25^.

While our structural, biochemical and simulation data indicate that *K. setae* ECR achieves high catalytic rates by synchronizing active sites, this might not necessarily be true for other ECRs. A differentiation into dimers of dimers was not observed in NADPH-bound or ternary structures of other ECRs (e.g., PDB: 4Y0K and 4A0S, respectively, which share substantial amino acid identity) (**SI Figure S11**). Note, however, that ECRs fall into two different classes. Primary ECRs (such as *K. setae* ECR) that operate in central carbon metabolism, and secondary ECRs (such as 4Y0K) that provide extender units for synthesizing polyketides in secondary metabolism. Whereas primary ECRs are under intense evolutionary pressure and show on average *k_cat_* values of 28 s^-1 26^, secondary ECRs are not selected for high catalytic rates, which is also reflected by one order of magnitude smaller average *k_cat_* value (*k_cat_* =1.2 s^-1^) ^26^. Thus, it might be tempting to speculate that secondary ECRs are not selected for high turnover rates during catalysis and, therefore, might not exhibit synchronized “half-site reactivity”, which is further supported by the observation that the residues Glu151, Asn218, and Asn157, conferring synchronization at the complex interface are not conserved in secondary ECRs (Figure 5c).

Altogether, our work revealed the structural-dynamics determinants explaining the substantial rate enhancement achieved by enoyl-CoA carboxylase/reductase through coordinated catalytic domain motions compared to unsynchronized catalytic reactions (e.g., in the four monomers of the triple variant E151D/N157E/N218E). The X-ray structures of the apo-form, binary and ternary ECR complexes provide the end states of these coordinated catalytic domain motions. PCA analysis of extensive molecular dynamics simulations delineates an atomistic picture of the coupled motions, with one opening and closing motion of the active site and a twist motion that might aid the substrate binding and product release. Further analysis shows that negative cooperativity and half-site reactivity are the governing principles of the substantial enhancement of overall reaction rate. Active site synchronization can be traced down to individual amino acids at the complex interface and the adenine binding pocket of the CoA substrate. These residues associated with the coupling of the catalytic domains across the different dimers are conserved among highly efficient ECRs from primary metabolism, enhancing overall catalytic rates by more than a factor of 30 compared to their counterparts in secondary metabolism.

## Methods

Details of experimental and computational methods are described in the extended supporting information. Briefly, these contain methods on amplification and cloning of K. setae ECR, site-directed mutagenesis of its residues at the subunit interface, adenine binding residues and intra-dimer communication residues, as well as cell lysis, protein purification, characterization of the oligomeric state of the enzyme, and spectrophotometric enzyme assays. Chemical synthesis of CoA-esters and analysis of ethylmalonyl-CoA stability are also described. Finally, we include detailed methods on crystallization of K.setae ECR complexes, data collection, processing and structure determination, QM/MM simulations of NADPH and substrate in the open and closed subunits of the ECR dimer, and molecular dynamics simulations of the conformational changes in the ECR tetramer.

## Data Availability

Coordinates of the four ECR structures have been deposited in the Protein Data Bank under accession codes, 6NA3 (apo), 6NA4 (Butyryl-CoA/NADPH bound), 6NA5 (NADP^+^ bound), and 6NA6 (NADPH-bound).

## Acknowledgments

The authors acknowledge Takanori Nakane from University of Tokyo for his help with calibration of XFEL data from SACLA, RIKEN, Japan. We would like to thank Eriko Nango, Rie Tanaka and RIKEN SPring-8 Center for their help with data collection at SACLA, RIKEN, Japan. H.D. acknowledges support from NSF Science and Technology Center grant NSF-1231306 (Biology with X-ray Lasers, BioXFEL). The XFEL experiments were performed at BL3 of SACLA with the approval of the Japan Synchrotron Radiation Research Institute (JASRI) (Proposal No. 2017A8055). The authors thank the beamline staff of Structural Molecular Biology Group, SSRL, SLAC and GM/CA CAT, Advance Photon Source, ANL for assistance on data collection. Y.R., R.G.S., M.S.H., and B.H. were supported by the U.S. Department of Energy, Office of Science, Office of Basic Energy Sciences under Contract No. DE-AC02-76SF00515. TJE and GS received support from the Max Planck Society, the European Research Council (ERC 637675 ‘SYBORG’), and the U.S. Department of Energy Joint Genome Institute a DOE Office of Science User Facility under Contract No. DE-AC02-05CH11231. D.A.S., A.G., H.G. and EVM thank the Max-Planck Society for funding as a Max-Planck-Partner group and CONICYT PCI MPG190003. A.G. thanks to CONICYT Doctorado Nacional grant 21190262. D.A.S. thanks Fondo Nacional de Desarrollo Científico y Tecnológico (Fondecyt) for his postdoctoral fellowship No. 3190579. H.D., Y.R., and S.W. are supported by DOE Office of Science, Biological Environmental Research, and National Institute of Health, NIGMS. C.G. and S.W. were supported by National Science Foundation, Major Research Instrument grant. H.D. and S.W. were partially supported by Stanford PRECOURT Institute. Gregory M. Stewart of SLAC and Moe Wakatsuki for graphics work.

